# Interplay between the human gut microbiome and host metabolism

**DOI:** 10.1101/561787

**Authors:** Alessia Visconti, Caroline I. Le Roy, Fabio Rosa, Niccolo Rossi, Tiphaine C. Martin, Robert P. Mohney, Weizhong Li, Emanuele de Rinaldis, Jordana T. Bell, J. Craig Venter, Karen E. Nelson, Tim D. Spector, Mario Falchi

**Author notes:** These authors contributed equally (Alessia Visconti, Caroline I. Le Roy). These authors share senior authorship (Tim D. Spector, Mario Falchi).

## Abstract

The human gut is inhabited by a complex and metabolically active microbial ecosystem regulating host health. While many studies have focused on the effect of individual microbial taxa, the metabolic potential of the entire gut microbial ecosystem has been largely under-explored. We characterised the gut microbiome of 1,004 twins via whole shotgun metagenomic sequencing (average 39M reads per sample). We observed greater similarity, across unrelated individuals, for functional metabolic pathways (82%) than for taxonomic composition (43%). We conducted a microbiota-wide association study linking both taxonomic information and microbial metabolic pathways with 673 blood and 713 faecal metabolites (Metabolon, Inc.). Metabolic pathways associated with 34% of blood and 95% of faecal metabolites, with over 18,000 significant associations, while species-level results identified less than 3,000 associations, suggesting that coordinated action of multiple taxa is required to affect the metabolome. Finally, we estimated that the microbiome mediated a crosstalk between 71% of faecal and 15% of blood metabolites, highlighting six key species (unclassified *Subdoligranulum* spp., *Faecalibacterium prausnitzii, Roseburia inulinivorans, Methanobrevibacter smithii, Eubacterium rectale*, and *Akkermansia muciniphila*). Because of the large inter-person variability in microbiome composition, our results underline the importance of studying gut microbial metabolic pathways rather than focusing purely on taxonomy to find therapeutic and diagnostic targets.

## Introduction

The human gut is home to trillions of microbes that form a complex community referred to as the gut microbiota. The metabolic activity of the gut microbiota is essential in maintaining host homeostasis and health, as proven, for instance, by the study of germfree animals [1,2]. Although the presence of a microbiota is vital, variations in its composition induces metabolic shifts that may result in alterations of host phenotype [3]. The gut microbiome is highly malleable and can be altered throughout lifespan mostly by environmental factors, such as diet and medication [4,5,6]. Although the external environment plays an important role in shaping the gut microbiome community, the host can affect the microbial ecosystem through its immune system, and has also a metabolic impact on the gut lumen [7,8,9].

The joint study of microbiome and metabolome has been suggested as the most promising approach to evaluate host-microbiome interactions [10]. However, studying the metabolic holobiont is complex, and few studies have tackled this issue in humans at any scale. Our group previously used 16S amplicon data to confirm that the gut microbiome is exceptionally metabolically active, and that the faecal metabolome may improve our estimation of the gut microbiota impact on health [11]. However, it is not possible to fully capture the metabolic activity of the gut microbiome using amplicon sequencing techniques alone, and the use of the more comprehensive whole shotgun metagenomic sequencing (WMGS) is necessary. Indeed, WMGS not only detects the taxonomic composition at higher resolution but also allows inferring its function, thus allowing the study of the metabolic potential of the microbial community. Here, we studied, in over a thousand twins, the effect of this metabolic activity on hosts health. We assessed the impact of the gut microbiome on both the gut and host systemic metabolism by using WMGS and untargeted faecal and blood metabolomics data. We found multiple associations between the gut microbiome (taxonomic composition and microbial metabolic function) and faecal and blood metabolites. In addition, we identified a number of microbial species and metabolic functions likely to play a leading role in the gut-systemic metabolic crosstalk.

## Results

### Gut microbiota composition is host-specific whereas its functions are shared across subjects

Whole metagenomic shotgun sequencing (WMGS) was performed on faecal samples provided by 1,073 volunteers from the TwinsUK registry, of which 1,004 surpassed quality control with an average of 39M high-quality microbial reads per sample (**Methods**, **Supplementary Table 1**). Taxonomic profiling identified, in the kingdoms of archaea and bacteria, 14 phyla, 24 classes, 37 orders, 74 families, 182 genera, and 580 species present in at least one sample (**Methods**). Each species was observed in a median of 2.7% of the samples, and 12% of species were sample-specific (**Figure 1**). The most ubiquitous species were from the *Subdoligranulum* genera (unclassified species), *Ruminococcus obeum, Ruminococcus torques*, and *Faecalibacterium prausnitzii*, all detected in more than 98% of the samples (**Supplementary Figure 1**). Microbial metabolic profiling (as described by the MetaCyc microbial metabolic pathways) identified 434 non-redundant pathways, which were detected in most samples (**Methods**). Each pathway was observed in a median of 91.6% of the samples, with 12% of the pathways present in all samples and only 2% being sample-specific (**Figure 1**).

**Table 1.** Association between species and the same named metabolite in faeces and blood. For each association, we report the number of observations (N), effect size (*β*), standard error (SE), and P value (P).

| Species | Metabolite | Faeces |  |  |  | Blood |  |  |  |
| --- | --- | --- | --- | --- | --- | --- | --- | --- | --- |
| | | N | $\beta$ | SE | P | N | $\beta$ | SE | P |
| *Akkermansia muciniphila* | p-cresol sulfate | 443 | -0.35 | 0.11 | $7.74 \times 10^{-4}$ | 829 | 0.47 | 0.10 | $6.36 \times 10^{-6}$ |
| *Bacteroidales bacterium ph8* | sebacate | 406 | 0.68 | 0.18 | $1.65 \times 10^{-4}$ | 718 | -0.50 | 0.09 | $4.33 \times 10^{-9}$ |
| *Eubacterium rectale* | p-cresol sulfate | 444 | 0.45 | 0.11 | $2.84 \times 10^{-5}$ | 823 | -0.45 | 0.11 | $2.09 \times 10^{-5}$ |
| *Methanobrevibacter smithii* | threonate | 287 | -0.65 | 0.14 | $3.31 \times 10^{-6}$ | 527 | 1.01 | 0.16 | $4.09 \times 10^{-10}$ |
| *Oscillibacter spp.* | 3-phenylpropionate | 418 | -0.54 | 0.11 | $1.95 \times 10^{-6}$ | 799 | -0.55 | 0.10 | $1.02 \times 10^{-7}$ |
| *Roseburia inulinivorans* | p-cresol sulfate | 433 | 0.43 | 0.10 | $1.09 \times 10^{-5}$ | 802 | -0.53 | 0.10 | $1.45 \times 10^{-7}$ |
| *Subdoligranulum spp.* | p-cresol sulfate | 461 | -0.32 | 0.10 | $9.86 \times 10^{-4}$ | 854 | 0.62 | 0.10 | $1.75 \times 10^{-9}$ |

**Fig. 1.**
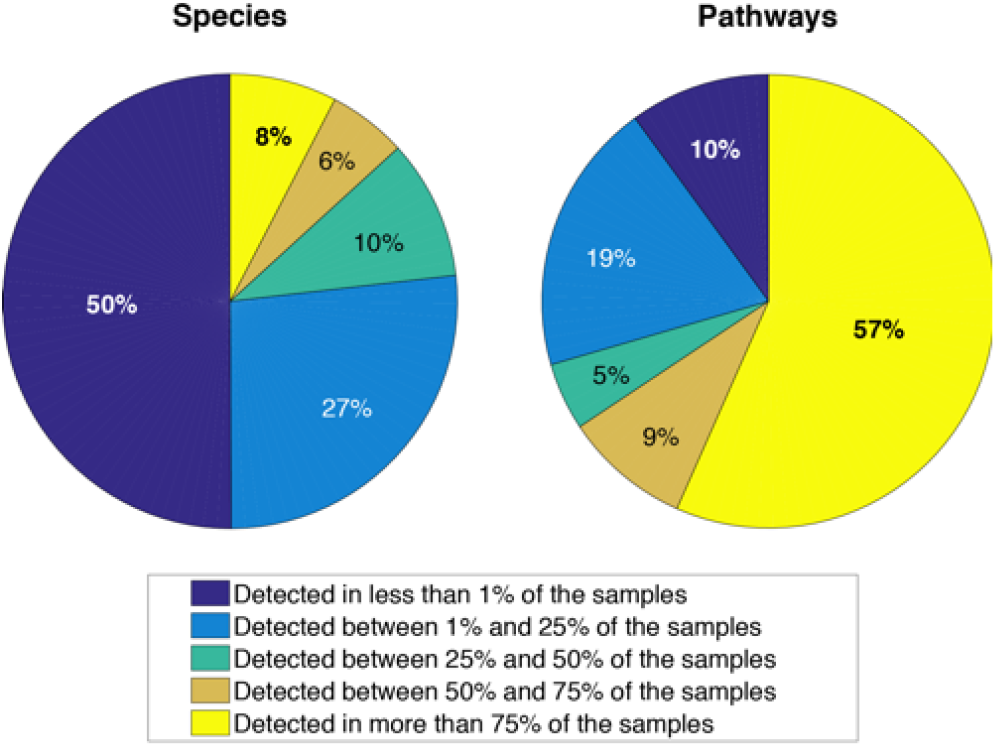
The composition of the gut ecosystem is unique to an individual while its functionality is maintained across the population. Pie charts representing the percentage of species (on the left) and pathways (on the right) present in less than 1% of the population (dark blue), between 1% and 25% (light blue), between 25% and 50% (turquoise), between 50% and 75% (brown), and more than 75% (yellow).

Microbial metabolic pathways were widely shared between individuals, compared to their taxonomical composition. Indeed, multiple known species (up to 465, and 29 on average) identified from the WMGS data, plus a large number of unclassified species, contributed to the abundance of each microbial metabolic pathway (**Supplementary Data D1**). As a consequence, pathway prevalence within our sample strongly correlated with the number of species in which it could be detected (Spearmans *ρ* = 0.34; P=9.4*×*10^-9^). When comparing unrelated individuals, we observed that, on average, they shared 82% of the pathways but only 43% of the species (paired Wilcoxon’s test P*<* 2*×*10^-16^; **Supplementary Figure 2**, **Methods**).

**Fig. 2.**
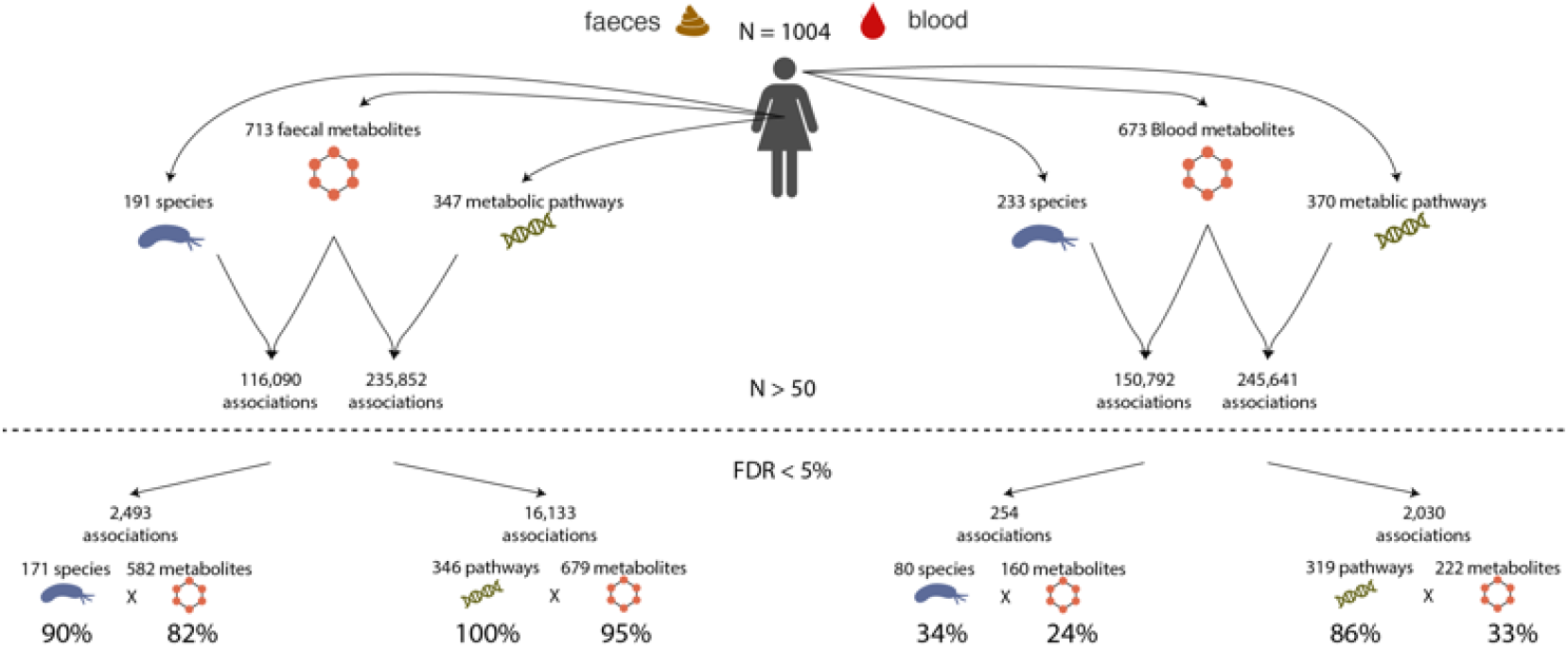
Study design and number of associations between the gut microbiome and the faecal and blood metabolome. The top of the Figure reports the number of microbial species and metabolic pathways which were detected in at least 50 individuals with metabolomics and WMGS data, and that were used in the study, and the number of associations tested. The bottom of the Figure reports the number of associations that were significant at a FDR 5%, along with the number and percentage of metabolites, microbial species, and microbial metabolic pathways involved.

### Microbiome composition and functions are strongly linked to gut lumen metabolism

Faecal metabolic profiles were available for 479 individuals with WMGS data. 713 annotated metabolites were measured in more than 50 individuals and tested for association with the gut microbiome at both taxonomic and functional levels (**Methods**, **Figure 2**). As expected, both the composition of the gut microbiome and its metabolic function were widely associated with the gut lumen metabolic content. At a 5% false discovery rate (FDR) we found 16,133 associations with microbial metabolic pathways and 2,493 associations with microbial species (**Supplementary Data D2 and D3**). In particular, 99.7% of the metabolic pathways were significantly associated with 95% of the faecal metabolites, while 90% of the species were associated with 82% of the faecal metabolites (**Methods; Figure 2**). On average, each metabolite level was associated with 4 species and 24 pathways. In addition, 145 (20%) metabolites were associated to a single species, while only 50 of them (7%) were associated to a single pathway. Five microbial species played a major metabolic role and were independently associated with 10% of the faecal metabolites (**Supplementary Figure 3**): unclassified *Subdoligranulum* spp. (149 metabolites), *Akkermansia muciniphila* (106 metabolites), *Roseburia inulinivorans* (105 metabolites), *Methanobrevibacter smithii* (96 metabolites), and *Roseburia intestinalis* (92 metabolites). In contrast, the top-five microbial metabolic pathways were associated with more than 53% of the faecal metabolites, with the pathways of L-rhamnose degradation I, Kdo transfer to lipid IVA III (Chlamydia), CDP diacylglycerol biosynthesis I and II and NAD biosynthesis I from aspartate affecting 226, 218, 215, 215, and 206 faecal metabolites, respectively.

**Fig. 3.**
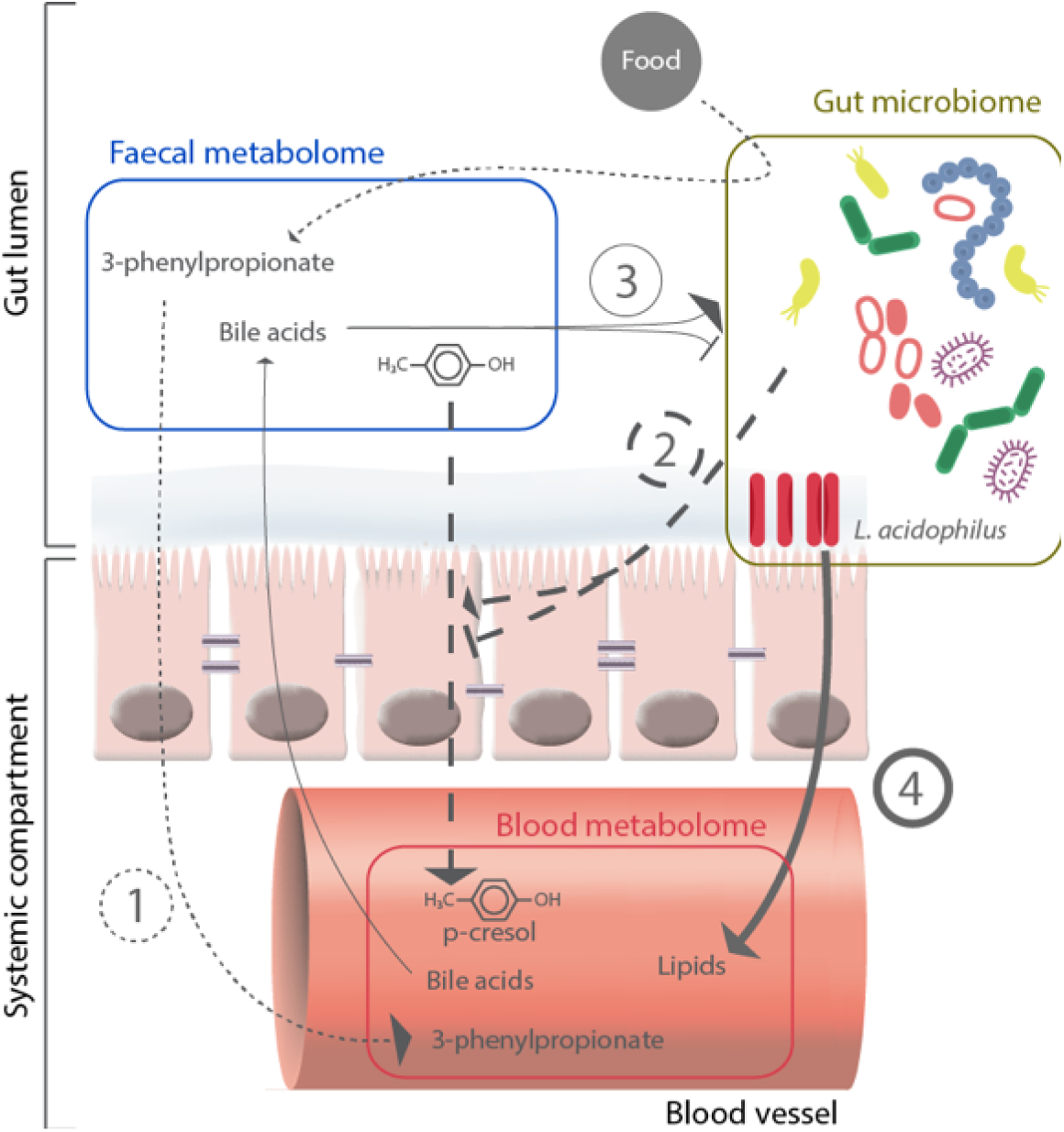
Summary of the possible mechanisms of host-microbiome metabolic crosstalk. The Figure highlights the possible four mechanisms implicated in the interactions between the gut microbiome, the faecal metabolome, and the blood metabolome. (1) Small dashed lines: metabolites are produced by the bacteria and then absorbed, resulting in associations between the microbiome and both the blood and faecal metabolites. (2) Large dashed lines: the microbiome affects the gut barrier integrity, resulting in alteration of metabolites absorption (i.e., the same metabolite is associated with a bacterium/pathway in both blood and faeces, but the directions of effects are opposite). (3) Light continuous line: metabolites produced by the host, such as bile acids, affect microbial growth. (4) Bold continuous line: direct bacteria to host cell interactions that results in host systemic modulation (i.e., bacteria are associated with blood metabolites but not with faecal metabolites).

We calculated the enrichment of the associated metabolites for metabolic super-pathways (as annotated by Metabolon, Inc.; **Methods**). Faecal metabolites associated with microbial species were enriched for a decrease in amino acids (adj P=1.6 *×*10^-4^) and an increase in lipids (adj P=1.9*×*10^-3^), while metabolites associated with metabolic pathways were enriched for a decrease in lipids (adj P=8.0*×*10^-5^), and an increase in both nucleotides (adj P=0.02) and carbohydrates (adj P=0.03).

B vitamins were strongly associated with both species and metabolic pathways, with riboflavin (vitamin B2), nicotinate (vitamin B3), pantothenate (vitamin B5), pyridoxine (vitamin B6), biotin (vitamin B7) associated with 9 to 27 species and with 48 to 155 microbial pathways (**Supplementary Data D2 and D3**). Finally, 16 associations were observed between vitamin E (alpha, beta, gamma and delta tocopherol) and species/pathways.

### Microbiome composition and functions associate with host systemic metabolites

Blood metabolomics profiling was available for 859 individuals with WMGS data. 673 annotated metabolites (including 369 metabolites also measured in faeces) were measured in more than 50 individuals, and used in this study. At a 5% FDR, we identified 2,030 associations with microbial metabolic pathways and 254 associations with microbial species (**Figure 2; Supplementary Data D4 and D5**). In particular, 86% and 34% of the microbial metabolic pathways and species, respectively, associated with 33% and 24% of the studied blood metabolites, respectively, with a total of 309 unique blood metabolites (46%) associated with the microbiome. The species showing the largest number of associations with blood metabolites were *Lactobacillus acidophilus* (N=30), and *Lactobacillus fermentum* (N=14; **Supplementary Figure 4**). The metabolite sebacate showed the highest number of associated species (N=11), followed by tartronate (N=9), phenylacetylglutamine (N=8), and p-cresol sulfate (N=6). On average, each blood metabolite was associated with two species (with 118 species controlling a single metabolite) and 10 metabolic pathways (with 93 pathways controlling a single metabolite). The three microbial metabolic pathways showing the largest number of associations with blood metabolites were the super pathways of L-phenylalanine and L-alanine biosynthesis, and the pathway of urate biosynthesis/inosine 5’-phosphate degradation (30, 26, and 24, respectively). Four blood metabolites were associated with more than 100 microbial metabolic pathways: phenylacetylglutamine (N=143) and p-cresol-glucuronide (N=102), two known gut microbial-derived metabolites, as well as tyramine O-sulfate (N=130), that can be synthesized by a *Eubacterium* enzyme [12], and 1,5-anhydroglucitol (N=129), that is present in a wide variety of food products.

### Microbial metabolic pathways have a broader metabolic footprint than species

Overall, we identified about seven times more associations between faecal and blood metabolites and microbial metabolic pathways than microbial species. We observed that pathways found in a larger number of species have a stronger impact on the metabolome, with a significant positive correlation between the number of species contributing to each pathway and the number of associations between the pathway and both faecal and blood metabolites (Spearmans *ρ*=0.27, P=2 *×*10^-6^, and *ρ*=0.33, P=1*×*10^-9^, respectively).

Our results confirmed a wide network of associations between the gut microbiome and the gut lumen metabolism, which extends to the systemic metabolome. At a 5% FDR, we identified 360 microbial metabolic pathways associating with 679 faecal and 222 blood metabolites, and 233 microbial species associating with 582 faecal and 160 blood metabolites. Most microbial metabolic pathways (85%) associating with one or more metabolites in the gut lumen also associated with one or more metabolites in blood (**Supplementary Table 2**). In contrast, the majority of microbial species only associated with faecal metabolites alone (58%, **Supplementary Table 2**). Still, 31% of species showed association with both faecal and blood metabolites, suggesting important effects on host systemic metabolism for this subset. Specifically, 4,861 pairs of faecal-blood metabolites co-associated with the same species and 108,565 pairs with the same metabolic pathway. Among these associations, 152 pairs involved exactly the same named metabolite in both faeces and blood (145 with metabolic pathways, and only seven with species; **Table 1**, **Supplementary Data D6**), while 113,274 pairs involved a different metabolite in faeces and blood (unique pairs N=27,608). Sebacate, threonate, and p-cresol sulfate, in both faeces and blood, showed the largest number of associations with pathways and with species in both faeces and blood.

### The microbiome is involved in the crosstalk between the gut lumen and host systemic metabolism

We further investigated the full set of faecal-blood co-associating pairs of metabolites to better understand whether the observed associations were randomly coincident at the same species or pathway, or if they were suggesting interplay between the gut and systemic environments. We assessed, through simulations, the probability that the microbiota was involved in the crosstalk between faecal and blood metabolites (**Methods**). We hypothesised that, if the species (or metabolic pathway) was involved in the crosstalk between faecal and blood metabolites, these were expected to more strongly correlate in individuals for which the species (or metabolic pathway) was present than in the remaining samples. We assessed this probability through simulations (**Methods**). Significantly higher correlations between co-associated faecal-blood metabolite pairs when species (or metabolic pathway) was detected (*P*_Empirical_ = 1*×*10^-3^ for species, and *P*_Empirical_ = 0.03 for pathways), suggesting extensive crosstalk between the gut and the systemic environments.

We further tested the presence of a faecal-blood crosstalk by applying the P-gain statistic to the co-associating faecal-blood metabolite pairs [13]. The P-gain statistic compares the increase in strength of association with the species (or pathway) when using the metabolite ratios compared to the smaller of the two p-values when using the two metabolite concentrations individually. A strong reduction in p-value indicates that two metabolites may be linked by a mechanism that involves the gut microbiota. To carefully assess a significance threshold for the P-gain statistics in our dataset, we estimated its empirical null distribution through simulations. We obtained a P-gain threshold of 73 for species and of 42 for metabolic pathways at an experimental-wide *α*-level of 0.05 (**Methods**). P-gains passing these thresholds are reported in **Supplementary Data D7 and D8**, and included 31% of the P-gains with species (1,325/4,232 co-associated metabolite pairs) and 19% with microbial metabolic pathways (16,839/88,452 co-associated metabolite pairs). The P-gain statistics suggested potential crosstalk between 36% of the faecal metabolites and 5% of the blood metabolites, involving 12% of the species (N=29). Crosstalk was wider with the microbial metabolic pathways, involving 70% of the faecal and 14% of blood metabolites, and 67% of the pathways (N=247).

At the species level, unclassified *Subdoligranulum* spp. accounted for 49% of the putative crosstalk, and *F. prausnitzii, R. inulinivorans, M. smithii, E. rectale*, and *A. muciniphila* together contributed to a further 36%. In contrast, the results at the pathway level were not dominated by a limited number of pathways, with the top six contributing only towards 24% of the observed crosstalk.

### Methanogens-host metabolic crosstalk is associated with adiposity

Threonate in blood showed the highest P-gains and a large number of significant faecal-blood interactions (with 61 faecal metabolites, including threonate levels in faeces). All interactions involved *M. smithii*, the main archeon in the human gut [14], present in 62% of our metagenomic samples. Threonate is produced from vitamin C under oxidative conditions [15]. In both blood and faeces, threonate was also associated with two microbial pathways linked to methanogenesis: coenzyme factor 420 biosynthesis (*β*=0.94, SE=0.18, P=2.2*×*10^-7^) and methanogenesis from H2 and CO2 (*β*=0.93, SE=0.18, P=3.3*×*10^-7^), to which *M. smithii* contributes, in our sample, for about 47% (the remaining attributable to *Methanosphaera stadtmanae, <* 1%, and to unclassified species, 53%; **Methods**). The role of *M. Smithii*, and of other methanogenic microbes in human health is still unclear, however, several studies suggested that its depletion is linked to obesity[16,17]. We found it to be significantly negatively associated with the percentage of visceral fat (*β*=-0.09, SE=0.04, P=0.013; **Supplementary Table 3**). We also observed a significant negative association (P¡0.05/3=0.017, **Supplementary Table 3**) between blood threonate and three measures of adiposity, namely BMI (*β*=-0.48, SE=0.12, P=1.2*×*10^-5^), and the percentage of total body (*β*=-0.41, SE=0.10, P=4.3*×*10^-5^) and visceral fat (*β*=-0.48, SE=0.11, P=2.6*×*10^-5^), while faecal threonate was not associated with any measure of adiposity (P¿0.05). Moreover, 31 out of 61 faecal metabolites whose crosstalk with blood threonate via *M. Smithii* was confirmed by the P-gain statistic were significantly associated with measures of adiposity (N=49, P*<* 0.05/(61 *×* 3) = 1.3 *×* 10^-3^; **Supplementary Data D9**).

## Discussion

Microbiome studies are mainly focused on the effect of individual microbial taxa on human health, while the metabolic potential of microbes has been largely overlooked. In line with recent observations [18], we confirmed that the microbiome metabolic potential (as described by the MetaCyc microbial metabolic pathways) is more conserved across unrelated subjects than species (82 vs 43%). We also observed that microbial metabolic pathways are highly redundant, with up to 465 identified species (and a possibly large number of unknown ones) sharing the same metabolic pathway.

Using 713 faecal and 673 blood metabolites measured by Metabolon, Inc. and WGMS data, we conducted a microbiota-wide association study. Our results showed that the gut metagenome (both at the species and at the metabolic pathway level) widely associates with both the gut and host systemic metabolism. The associations with faecal metabolites suggest that most microbial species (90%) and metabolic pathways (99.7%) interact with their surrounding metabolic environment. In particular, metabolic pathways were significantly associated with 95% of the faecal metabolites, while microbial species were associated with 82% of the faecal metabolites. The results for species were comparable to those previously reported in a recent study on the TwinsUK cohort leveraging 16S amplicon data [11]. In both studies, we observed that over 90% of microbes were associated with a vast proportion of the measured gut metabolites (*>*80%). However, the WMGS data used in this study improved the precision of the taxonomic results, allowing identification of associations at the species level. For instance, we were able to identify five species interacting with at least 10% of the studied faecal metabolites. Four of them (*Subdoligranulum spp., A. muciniphila, R. inulinivorans, and R. intestinalis*) were present in at least 80% of the population and are already known for their ability to affect gut lumen metabolism [19,20,21,22]. Interestingly, among the numerous microbiome-metabolome associations identified in this study, a large proportion was involved with the metabolism of vitamins. For instance, we observed over 700 associations with vitamin B-related metabolites. While B vitamins are mostly provided to the host through diet, these can also be synthesised by lactic acid bacteria [23]. Our results show a similar number of positive and negative associations with vitamin B metabolites, suggesting that the microbiome is not only involved in the biosynthesis of vitamins B but also in its degradation. In this study, we also evaluated the impact of the gut microbiome on the host systemic metabolism. We showed that nearly half of the blood metabolites (N=309, 46%) were associated with microbial species and/or metabolic pathways. More exactly, 34% of the species and 86% of the pathways were associated with 24% and 33% of the metabolites, respectively. Two bacteria stood out as playing a major role: *L. acidophilus* (5%) and *L. fermentum* (2%), both known for their probiotic properties [24,25,26].

Notably, a previous study on the TwinsUK cohort observed that 72% of blood metabolites were under host genetic influence [27]. Interestingly, 144 out of 309 microbiome-associated blood metabolites (47%) identified in our study were not heritable. Heritabilities for the remaining 165 blood metabolites ranged from 10% to 78%, with a mean value of 47% (**Supplementary Data D10**). This suggests that, despite the widespread host genetic effects on blood metabolites, the gut microbiome might play a role on the systemic metabolism that is independent from the host genome.

Altogether, our results indicate an intense interplay between the gut microbiome and its host. While only a small number of metabolites were found to be associated to a same species (or pathway) in both metabolic environments (N=152), we detected more than 27,000 unique pairs of faecal-blood metabolites which were associated with the same microbial species and/or metabolic pathway (co-associated metabolites). The limited size of our study sample makes it unsuitable to test causality using a Mendelian randomisation method [28]. Nonetheless, using two complementary approaches, we showed that, first, co-associated metabolites are more strongly correlated in the presence of the associated species or metabolic pathways (*P*_Empirical_ = 1*×*10^-3^ and 0.03, respectively), and, second, that a significant crosstalk, as assessed through the P-gain statistic, exists between 71% of the faecal and the 15% of blood metabolites, involving 12% of the species and 67% of the pathways. We suggest that four potential mechanisms could underlie the interplay between these two metabolomic environments (**Figure 3**). First, these interactions could be triggered by the metabolic activity of the microbiome [29]. Second, the gut microbiome could mediate metabolite transfer through the gut barrier by affecting its integrity, as suggested, for example, by the associations involving the same named species and metabolites in both blood and faeces (**Table 1**). Indeed, these associations showed opposite direction of effects, suggesting that microbes may modulate the absorption of the metabolites by the host rather than its bioavailability. Third, microbial growth could be impacted by secretion of metabolites by the host within the gut lumen as extensively discussed regarding bile acids [30,31]. Fourth, host-gut microbiome interactions could also be triggered by non-metabolic interactions including microbial secretion of peptides or direct cell-cell interactions [32], which could not be investigated in the present study.

We observed about seven times more associations between metabolites and microbial metabolic pathways than species. This trend was even stronger when studying the faecal-blood crosstalk, with nearly 13 times more co-associated metabolite pairs identified by means of the P-gain statistics for microbial metabolic pathways than species. These results support the claim that looking at functions rather than taxonomy alone gives a better appreciation of the true gut microbiome metabolic activity [10]. We suggest that the large number of associations with metabolic pathways could be mostly due to the fact that they are usually conserved among different species. Furthermore, the majority of the metabolic pathways are associated with metabolites apparently unrelated to their functions. Therefore, associations with a metabolic pathway could be interpreted also as associations with a particular microbial sub-community rather than only to its specific function.

This study has some limitations. First, we used data from a cohort including only individuals of European ancestry and composed predominantly of middle-aged woman (96%, average age 65 years old). Therefore, our results may not generalise to diverse populations. Ideally, data collected in other larger cohorts and meta-analyses would be necessary to confirm our novel findings. Second, despite the large-scale sample, this is a cross-sectional study, and no causal relationship between the microbiome and the metabolome can be inferred from the identified associations. Third, while WMGS data allow us to infer the functional capability of the microbial community, it does not provide information on which microbial metabolic pathways are actually active. Metatranscriptomic data will help in bridging this gap, also allowing discerning between associations with microbial metabolic pathways that are connected to their specific function or that are simply a proxy for microbial subcommunities.

In conclusion, we first confirmed the key role played by the microbiome on the gut lumen and host systemic metabolism. Next, we described the microbiome effect on the interplay between the two metabolic compartments. We observed that only a few key species, but many common microbial functions, are substantially associated with faecal and blood metabolic profiles. Therefore, microbial metabolic pathways should be considered beyond their primary function and interpreted as proxies for microbial communities, interacting with their surrounding environment. Future treatments designed to improve host health through the modulation of the gut microbiome should optimally target functionally-related microbial communities rather than single organisms.

## Methods

### DNA extraction, library preparation, and metagenomic sequencing

A 3-mL volume of lysis buffer (20 mM Tris-HCl pH 8.0, 2 mM Sodium EDTA 1.2% Triton X-100) was added to 0.5 g of stool sample, and the sample vortexed until homogenized. A 1.2 mL volume of homogenized sample and 15 mL of Proteinase K (Sigma Aldrich, PN.P2308) enzyme was aliquoted to a 1.5 mL tube with garnet beads (Mo Bio PN. 1283050-BT). Bead tubes were then incubated at 65^°^C for 10 min and then 95^°^C for 15 min. Tubes were then placed in a Vortex Genie 2 to perform bead beating for 15 min and the sample subsequently spun in an Eppendorf Centrifuge 5424. 800 uL of supernatant was then transferred to a deep well block and DNA extracted and purified using a Chemagic MSM I (Perkin Elmer) following the manufacturers protocol. Zymo Onestep Inhibitor Removal kit was then performed following manufacturers instructions (Zymo Research PN. D6035). DNA samples were then quantified using Quant-iT on an Eppendorf AF2200 plate reader.

Nextera XT libraries were prepared manually following the manufacturers protocol (Illumina, PN. 15031942). Briefly, samples were normalized to 0.2 ng/ml DNA material per library using a Quant-iT picogreen assay system (Life Technologies, PN. Q33120) on an AF2200 plate reader (Eppendorf), then fragmented and tagged via tagmentation. Amplification was performed by Veriti 96 well PCR (Applied Biosystems) followed by AMPure XP bead cleanup (Beckman Coulter, PN. A63880). Fragment size for all libraries were measured using a Labchip GX Touch HiSens. Sequencing was performed on an Illumina HiSeq 2500 using SBS kit V4 Chemistry, with a read length of 2×125 bp. Sequencing yielded an average number of paired-end reads of 26.73M.

### Taxonomic profiling, and functional annotation

Paired-end reads were processed using the YAMP pipeline [33]. Briefly, we first removed identical reads, potentially generated by PCR amplification [34]. Next, reads were filtered to remove adapters, known artefacts, phix174, and then quality trimmed (PhRED quality score *<* 10). Reads that became too short after trimming (N *<* 60 bp) were discarded. We retained singleton reads (*i.e.*, reads whose mate has been discarded) in order to retain as much information as possible. Contaminant reads belonging to the host genome were removed (build: GRCh37). Low quality samples, *i.e.*, samples with less than 15M reads after QC were discarded (N=4). Next, MetaPhlAn2 [35] (v. 2.6.0) and the HUMAnN2 pipeline[18] (v 0.10.0), both included into the YAMP pipeline, were used to characterise the microbial community composition and its functional capabilities, respectively. Functional capabilities of the microbial community were described by the MetaCyc metabolic pathways, and assessed using the UniRef90 proteomic database annotations. HUMAnN2 was also used to evaluate the percentage of species contributing to the abundance of each microbial metabolic pathway.

A principal component analysis evaluated using the taxonomic profiling was used to identify and discard ecologically abnormal samples (N=37). If sample scores were greater than 3 times the standard deviation on one of the first 10 principal components the sample was labelled as outlier and discarded. Finally, we removed individuals not of European ancestry (N=9, self-reported via questionnaire) resulting in 1,004 samples with an average number of reads of 39M (39 males, 965 female), all living in the UK at the time of specimen collection (**Supplementary Table 1**). The dataset included 161 monozygotic twin pairs (N=322), 201 dizygotic twin pairs (N=402), and 280 singletons.

Taxonomic and microbial metabolic pathways relative abundances were arcsine squareroot transformed, filtered for outliers using the Grubbs outlier test (significance threshold P=0.05), and standardised to have zero mean and unit variance [36]. Under the assumption that a zero relative abundance meant impossibility to detect the taxum/pathway rather than its absence, zero values were considered as not available (NA).

### Metabolomics profiling

Metabolite concentrations were measured from fecal samples and blood by Metabolon, Inc., Morrisville, North Carolina, USA, by using an untargeted LC/MS/MS platform as previously described [11,27]. A total of 1,116 metabolites were measured in the 480 faecal samples, including 850 of known chemical identity used in this study. In blood, a total of 902 metabolites were measured in 859 individuals, 687 of which had known chemical identities. Metabolites were scaled by run-day medians, and log-transformed. Faecal metabolites were further scaled to have mean zero and standard deviation one. Metabolites that were indicated as below detection level (zero) were considered as not available (NA).

### Within-unrelated individuals species and microbial metabolic pathways similarity

For all individuals in our datasets, we codified the absence/presence of a microbial species/ metabolic pathways with 0 and 1, respectively. Then, after having identified all possible pairs of unrelated individuals (N=1,006,288), we evaluated their similarity as the ratio between the number of species/pathways which were present in both members of the pair and the number of species/metabolic pathways which were present in at least one of the members of the pair. The similarity distributions were then compared using a paired Wilcoxon’s test.

### Metagenome-wide association study

Associations of faecal and blood metabolites with species and microbial metabolic pathways transformed relative abundances were carried out using PopPAnTe [37], which uses a variance component framework and the matrix of the expected kinship between each pair of individuals to model the resemblance among family members. Sex and age at the sample collection were included as covariates. Only pairs of metabolites-species/pathways with at least 50 observations were tested for association. The significance of the associations was evaluated by comparing the likelihood of a full model, including the species/metabolic pathways in the fixed effect, and the likelihood of a null model where these effects were constrained to zero. Associations passing a false discovery rate (FDR) threshold of 5% were considered significant. FDR was evaluated using Storeys method [38].

### Enrichment analysis

Enrichment analysis was performed using the super-pathways annotation provided by Metabolon, Inc., as described elsewhere [11]. Metabolites were grouped in the following eight super-pathways: amino acid, carbohydrate, cofactors and vitamins, energy, lipid, nucleotide, peptide, and xenobiotics. Enrichment P values were evaluated using the parametric analysis of gene set enrichment (PAGE) algorithm [39] using 10,000 random permutations as implemented in the *piano* R package [40] (v 1.20). The PAGE algorithm, being based on a two-tailed Z score, can evaluate whether each super-pathway is significantly enriched for an increase or a decrease of the amount of metabolites which it includes.

### Gut-host metabolic crosstalk

We selected all pairs of metabolites that were observed in at least 100 individuals and were associated with the same species/metabolic pathway in both environments (co-associated metabolites) and used two approaches to detect and validate the presence of an interplay between the gut and the systemic host metabolism. First, we hypothesised that, if a species (or pathway) is involved in the crosstalk between faecal and blood metabolites, these metabolites would be expected to be more strongly correlated in the presence of the species (or pathway) than in its absence. We used the missingness observed in our WMGS data to test this hypothesis. Indeed, while we are not able to measure it, we can confidently assume that a variable proportion of missing data in our dataset are likely to include truly missing species (or pathways). We tested this hypothesis through simulations. We selected all pairs of co-associated metabolites interacting with species (or pathways) with at least 30 missing observations, and built 1,000 random datasets which included 1,000 pairs of metabolites matched by correlation and sample size to the original set of co-associated metabolites. These new pairs were then combined with species (or pathways) having the same missingness pattern of the actual associated species (or pathways). Then, we used these simulated datasets to assess the probability of observing increased correlation between metabolites when the species (or pathway) was present in the co-associated metabolites compared to the matched pairs. Second, we evaluated, for each pair of co-associated metabolites, its Pgain statistic, which allows determining whether the ratio between the two metabolites is more informative than the single metabolites alone, therefore suggesting the presence of a relationship between them [13]. To this aim, first, we evaluated all the log ratios between each pair of co-associated metabolites. Then, we associated the obtained ratios with the specific species/pathway by fitting a linear mixed effect model in R (package *lme4*, version 1.1.18), including age and sex as fixed effects, and family structure as a random effect. Finally, we evaluated the P-gain statistic as the ratio between the minimum P value obtained using the single metabolites alone and the P value obtained using their ratio [13]. It has previously suggested that a critical P-gain threshold taking into account multiple test correction, under the assumption of a type I error rate of 0.05, would be 10 times the number of tests [13]. However, it has also been observed that the magnitude of the P-gain statistic can be reduced by the increasing correlation between the metabolites and their ratio, and increased by an increasing sample size [13], two parameters which varied greatly in our dataset. We, therefore, estimated a null distribution empirically using a conservative assumption of no interplay between metabolites associated with different species or pathways. Therefore, we build a null distribution of P-gain statistics using 100,000 pairs of randomly selected metabolites which were associated at 5% FDR with two different species (or pathways) but were matched one-to-one by correlation and sample size to the co-associated metabolite pairs. We used the top 5% P-gain value as the critical P-gain threshold.

### Adiposity phenotypes data and association study

Subjects were asked to remove their shoes, and height (in cm) was measured using a stadiometer. Weight (in kg) was measured on digital scales. Total and visceral fat mass percentage was determined in 1,141 individuals with metagenomic and/or metabolomic data available by DXA (Dual-Energy X-ray Absorptiometry; Hologic QDR; Hologic, Inc., Waltham, MA, USA) whole-body scanning by a trained research nurse, as described elsewhere [41]. The QDR System Software Version 12.6 (Hologic, Inc., Waltham, MA, USA) was used to analyse the scans. Measurements greater than 3 standard deviations from the dataset mean were excluded from the analysis. To ensure the normality of their distribution, the data were rank-based inverse normalized. Associations with *M. Smithii*, blood and faecal threonate, and 61 faecal metabolites whose crosstalk with blood threonate via M. smithii was confirmed by the P-gain statistic, were carried out by fitting a linear mixed effect model in R (package *lme4*, version 1.1.18), including age and sex as fixed effects, and family structure as a random effect.

## Competing interests

The following authors are or were employees of Human Longevity, Inc.: W.L., J.C.V., K.E.N. R.P.M is a current employee of Metabolome, Inc. E.d.R is a current employee of Sanofi. T.D.S. is a consultant for Zoe Global Ltd. The other authors declare that they have no conflict of interest.

## Acknowledgments

TwinsUK was funded by the Wellcome Trust and MRC. The study also receives support from the National Institute for Health Research (NIHR) funded BioResource, Clinical Research Facility and Biomedical Research Centre based at Guys and St Thomas NHS Foundation Trust in partnership with Kings College London. We gratefully acknowledge support provided by the JPI HDHL funded DINAMIC consortium (administered by the MRC UK, MR/N030125/1).

## References

1. S. P. Claus, T. M. Tsang, Y. Wang, O. Cloarec, E. Skordi, F.-P. Martin, S. Rezzi, A. Ross, R. Kochhar, E. Holmes, and J. K. Nicholson, “Systemic multicompartmental effects of the gut microbiome on mouse metabolic phenotypes,” Molecular Systems Biology, vol. 4, Oct. 2008.

2. W. R. Wikoff, A. T. Anfora, J. Liu, P. G. Schultz, S. A. Lesley, E. C. Peters, and G. Siuzdak, “Metabolomics analysis reveals large effects of gut microflora on mammalian blood metabolites,” Proceedings of the National Academy of Sciences, vol. 106, pp. 3698–3703, Mar. 2009.

3. P. J. Turnbaugh, R. E. Ley, M. A. Mahowald, V. Magrini, E. R. Mardis, and J. I. Gordon, “An obesity-associated gut microbiome with increased capacity for energy harvest,” Nature, vol. 444, pp. 1027–1031, Dec. 2006.

4. J. Goodrich, J. Waters, A. Poole, J. Sutter, O. Koren, R. Blekhman, M. Beaumont, W. VanTreuren, R. Knight, J. Bell, T. Spector, A. Clark, and R. Ley, “Human Genetics Shape the Gut Microbiome,” Cell, vol. 159, pp. 789–799, Nov. 2014.

5. J. Goodrich, E. Davenport, M. Beaumont, M. Jackson, R. Knight, C. Ober, T. Spector, J. Bell, A. Clark, and R. Ley, “Genetic Determinants of the Gut Microbiome in UK Twins,” Cell Host & Microbe, vol. 19, pp. 731–743, May 2016.

6. H. Xie, R. Guo, H. Zhong, Q. Feng, Z. Lan, B. Qin, K. J. Ward, M. A. Jackson, Y. Xia, X. Chen, B. Chen, H. Xia, C. Xu, F. Li, X. Xu, J. Y. Al-Aama, H. Yang, J. Wang, K. Kristiansen, J. Wang, C. J. Steves, J. T. Bell, J. Li, T. D. Spector, and H. Jia, “Shotgun Metagenomics of 250 Adult Twins Reveals Genetic and Environmental Impacts on the Gut Microbiome,” Cell Systems, vol. 3, pp. 572–584.e3, Dec. 2016.

7. J. K. Nicholson, E. Holmes, J. Kinross, R. Burcelin, G. Gibson, W. Jia, and S. Pettersson, “Host-Gut Microbiota Metabolic Interactions,” Science, vol. 336, pp. 1262–1267, June 2012.

8. L. V. Hooper, D. R. Littman, and A. J. Macpherson, “Interactions Between the Microbiota and the Immune System,” Science, vol. 336, pp. 1268–1273, June 2012.

9. A. M. Valdes, J. Walter, E. Segal, and T. D. Spector, “Role of the gut microbiota in nutrition and health,” BMJ, p. k2179, June 2018.

10. P. J. Turnbaugh and J. I. Gordon, “An Invitation to the Marriage of Metagenomics and Metabolomics,” Cell, vol. 134, pp. 708–713, Sept. 2008.

11. J. Zierer, M. A. Jackson, G. Kastenmller, M. Mangino, T. Long, A. Telenti, R. P. Mohney, K. S. Small, J. T. Bell, C. J. Steves, A. M. Valdes, T. D. Spector, and C. Menni, “The fecal metabolome as a functional readout of the gut microbiome,” Nature Genetics, vol. 50, pp. 790–795, June 2018.

12. K. Kobashi, Y. Fukaya, D. H. Kim, T. Akao, and S. Takebe, “A novel type of aryl sulfotransferase obtained from an anaerobic bacterium of human intestine,” Archives of Biochemistry and Biophysics, vol. 245, pp. 537–539, Mar. 1986.

13. A.-K. Petersen, J. Krumsiek, B. Wgele, F. J. Theis, H.-E. Wichmann, C. Gieger, and K. Suhre, “On the hypothesis-free testing of metabolite ratios in genome-wide and metabolome-wide association studies,” BMC Bioinformatics, vol. 13, no. 1, p. 120, 2012.

14. E. E. Hansen, C. A. Lozupone, F. E. Rey, M. Wu, J. L. Guruge, A. Narra, J. Goodfellow, J. R. Zaneveld, D. T. McDonald, J. A. Goodrich, A. C. Heath, R. Knight, and J. I. Gordon, “Pangenome of the dominant human gut-associated archaeon, Methanobrevibacter smithii, studied in twins,” Proceedings of the National Academy of Sciences, vol. 108, pp. 4599–4606, Mar. 2011.

15. G. L. Simpson and B. J. Ortwerth, “The non-oxidative degradation of ascorbic acid at physiological conditions,” Biochimica Et Biophysica Acta, vol. 1501, pp. 12–24, Apr. 2000.

16. M. Million, M. Maraninchi, M. Henry, F. Armougom, H. Richet, P. Carrieri, R. Valero, D. Rac- cah, B. Vialettes, and D. Raoult, “Obesity-associated gut microbiota is enriched in Lactobacillus reuteri and depleted in Bifidobacterium animalis and Methanobrevibacter smithii,” International Journal of Obesity, vol. 36, pp. 817–825, June 2012.

17. A. Schwiertz, D. Taras, K. Schfer, S. Beijer, N. A. Bos, C. Donus, and P. D. Hardt, “Microbiota and SCFA in Lean and Overweight Healthy Subjects,” Obesity, vol. 18, pp. 190–195, Jan. 2010.

18. E. A. Franzosa, L. J. McIver, G. Rahnavard, L. R. Thompson, M. Schirmer, G. Weingart, K. S. Lipson, R. Knight, J. G. Caporaso, N. Segata, and C. Huttenhower, “Species-level functional profiling of metagenomes and metatranscriptomes,” Nature Methods, vol. 15, pp. 962–968, Nov. 2018.

19. M. Derrien, “Akkermansia muciniphila gen. nov., sp. nov., a human intestinal mucin-degrading bacterium,” INTERNATIONAL JOURNAL OF SYSTEMATIC AND EVOLUTIONARY MICROBIOLOGY, vol. 54, pp. 1469–1476, Sept. 2004.

20. S. H. Duncan, A. Barcenilla, C. S. Stewart, S. E. Pryde, and H. J. Flint, “Acetate Utilization and Butyryl Coenzyme A (CoA):Acetate-CoA Transferase in Butyrate-Producing Bacteria from the Human Large Intestine,” Applied and Environmental Microbiology, vol. 68, pp. 5186–5190, Oct. 2002.

21. B. S. Samuel and J. I. Gordon, “A humanized gnotobiotic mouse model of host-archaeal-bacterial mutualism,” Proceedings of the National Academy of Sciences, vol. 103, pp. 10011–10016, June 2006.

22. B. S. Samuel, E. E. Hansen, J. K. Manchester, P. M. Coutinho, B. Henrissat, R. Fulton, P. Latreille, K. Kim, R. K. Wilson, and J. I. Gordon, “Genomic and metabolic adaptations of Methanobrevibacter smithii to the human gut,” Proceedings of the National Academy of Sciences, vol. 104, pp. 10643–10648, June 2007.

23. J. G. LeBlanc, C. Milani, G. S. de Giori, F. Sesma, D. van Sinderen, and M. Ventura, “Bacteria as vitamin suppliers to their host: a gut microbiota perspective,” Current Opinion in Biotechnology, vol. 24, pp. 160–168, Apr. 2013.

24. K. Kailasapathy and J. Chin, “Survival and therapeutic potential of probiotic organisms with reference to *Lactobacillus acidophilus* and *Bifidobacterium* spp.,” Immunology and Cell Biology, vol. 78, pp. 80–88, Feb. 2000.

25. Y. Bao, Y. Zhang, Y. Zhang, Y. Liu, S. Wang, X. Dong, Y. Wang, and H. Zhang, “Screening of potential probiotic properties of Lactobacillus fermentum isolated from traditional dairy products,” Food Control, vol. 21, pp. 695–701, May 2010.

26. J. Maldonado, F. Caabate, L. Sempere, F. Vela, A. R. Snchez, E. Narbona, E. Lpez-Huertas, A. Geerlings, A. D. Valero, M. Olivares, and F. Lara-Villoslada, “Human Milk Probiotic Lactobacillus fermentum CECT5716 Reduces the Incidence of Gastrointestinal and Upper Respiratory Tract Infections in Infants:,” Journal of Pediatric Gastroenterology and Nutrition, vol. 54, pp. 55–61, Jan. 2012.

27. T. Long, M. Hicks, H.-C. Yu, W. H. Biggs, E. F. Kirkness, C. Menni, J. Zierer, K. S. Small, M. Mangino, H. Messier, S. Brewerton, Y. Turpaz, B. A. Perkins, A. M. Evans, L. A. D. Miller, L. Guo, C. T. Caskey, N. J. Schork, C. Garner, T. D. Spector, J. C. Venter, and A. Telenti, “Whole-genome sequencing identifies common-to-rare variants associated with human blood metabolites,” Nature Genetics, vol. 49, pp. 568–578, Apr. 2017.

28. G. Davey Smith and S. Ebrahim, “Mendelian randomization: can genetic epidemiology contribute to understanding environmental determinants of disease?*,” International Journal of Epidemiology, vol. 32, pp. 1–22, Feb. 2003.

29. B. K. I. Meijers and P. Evenepoel, “The gut-kidney axis: indoxyl sulfate, p-cresyl sulfate and CKD progression,” Nephrology Dialysis Transplantation, vol. 26, pp. 759–761, Mar. 2011.

30. J. R. Swann, E. J. Want, F. M. Geier, K. Spagou, I. D. Wilson, J. E. Sidaway, J. K. Nicholson, and E. Holmes, “Systemic gut microbial modulation of bile acid metabolism in host tissue compartments,” Proceedings of the National Academy of Sciences, vol. 108, pp. 4523–4530, Mar. 2011.

31. J. M. Ridlon, D. J. Kang, P. B. Hylemon, and J. S. Bajaj, “Bile acids and the gut microbiome:,” Current Opinion in Gastroenterology, vol. 30, pp. 332–338, May 2014.

32. J. M. Wells, O. Rossi, M. Meijerink, and P. van Baarlen, “Epithelial crosstalk at the microbiota-mucosal interface,” Proceedings of the National Academy of Sciences, vol. 108, pp. 4607–4614, Mar. 2011.

33. A. Visconti, T. C. Martin, and M. Falchi, “YAMP: a containerized workflow enabling reproducibility in metagenomics research,” GigaScience, vol. 7, July 2018.

34. M. B. Jones, S. K. Highlander, E. L. Anderson, W. Li, M. Dayrit, N. Klitgord, M. M. Fabani, V. Seguritan, J. Green, D. T. Pride, S. Yooseph, W. Biggs, K. E. Nelson, and J. C. Venter, “Library preparation methodology can influence genomic and functional predictions in human microbiome research,” Proceedings of the National Academy of Sciences, vol. 112, no. 45, pp. 14024–14029, 2015.

35. D. T. Truong, E. A. Franzosa, T. L. Tickle, M. Scholz, G. Weingart, E. Pasolli, A. Tett, C. Huttenhower, and N. Segata, “MetaPhlAn2 for enhanced metagenomic taxonomic profiling,” Nature Methods, vol. 12, p. 902, Sept. 2015.

36. J. Lloyd-Price, A. Mahurkar, G. Rahnavard, J. Crabtree, J. Orvis, A. B. Hall, A. Brady, H. H. Creasy, C. McCracken, M. G. Giglio, D. McDonald, E. A. Franzosa, R. Knight, O. White, and C. Huttenhower, “Strains, functions and dynamics in the expanded Human Microbiome Project,” Nature, Sept. 2017.

37. A. Visconti, M. Al-Shafai, W. A. Al Muftah, S. B. Zaghlool, M. Mangino, K. Suhre, and M. Falchi, “PopPAnTe: population and pedigree association testing for quantitative data,” BMC Genomics, vol. 18, Dec. 2017.

38. J. D. Storey, “A direct approach to false discovery rates,” Journal of the Royal Statistical Society: Series B (Statistical Methodology), vol. 64, pp. 479–498, Aug. 2002.

39. S.-Y. Kim and D. J. Volsky, “PAGE: parametric analysis of gene set enrichment,” BMC bioinformatics, vol. 6, p. 144, June 2005.

40. L. Vremo, J. Nielsen, and I. Nookaew, “Enriching the gene set analysis of genome-wide data by incorporating directionality of gene expression and combining statistical hypotheses and methods,” Nucleic Acids Research, vol. 41, pp. 4378–4391, Apr. 2013.

41. A. Moayyeri, C. J. Hammond, D. J. Hart, and T. D. Spector, “Effects of age on genetic influence on bone loss over 17 years in women: The Healthy Ageing Twin Study (HATS),” Journal of Bone and Mineral Research, vol. 27, pp. 2170–2178, Oct. 2012.

