## Supplementary Material for "Interplay between the human gut microbiome and host metabolism"

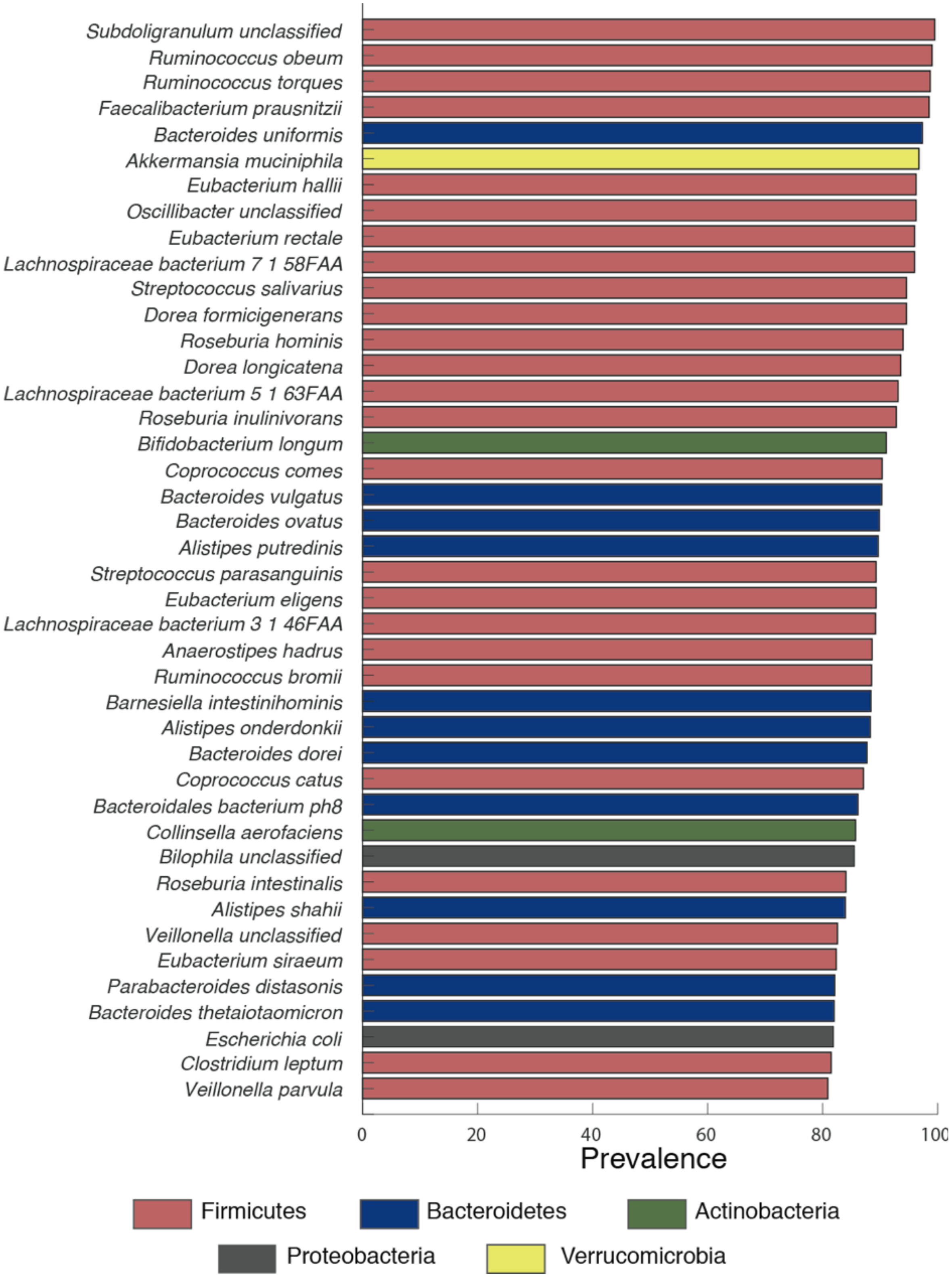


Supplementary Figure S1: Species detected in at least 80% of the population.

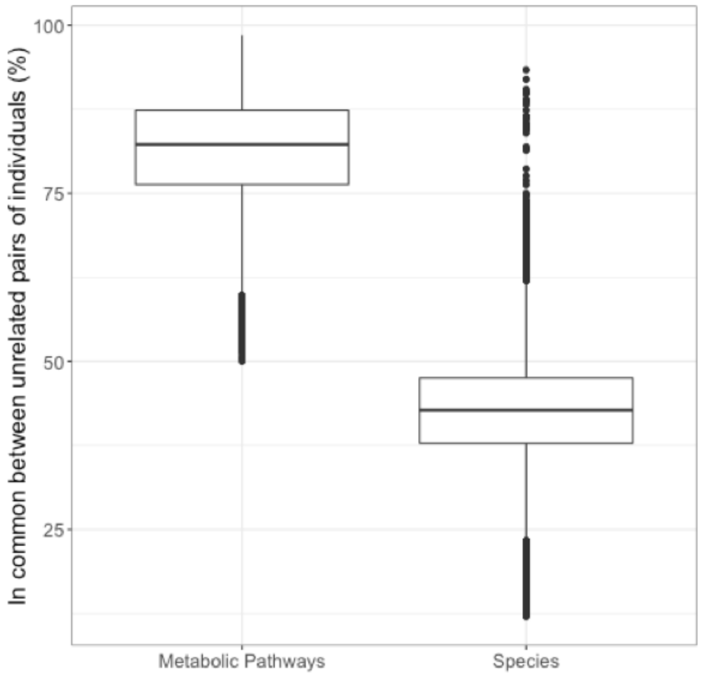


Supplementary Figure S2: Percentage of microbial metabolic pathways and species in common between pairs of unrelated individuals. A pathway/species was considered in common when measured in both individuals, while pathways/species measured in only one of them was considered individual-specific. Pathways/species not measured in either individual were not considered.

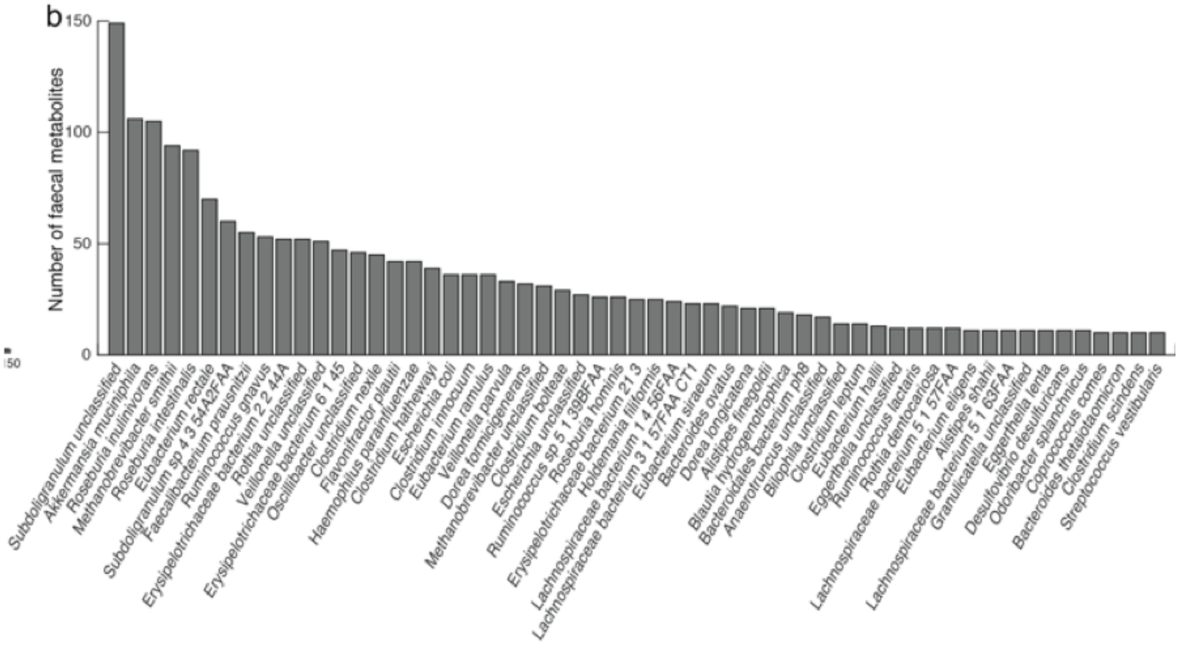


Supplementary Figure S3: Number of faecal metabolites significantly associated with microbial species.

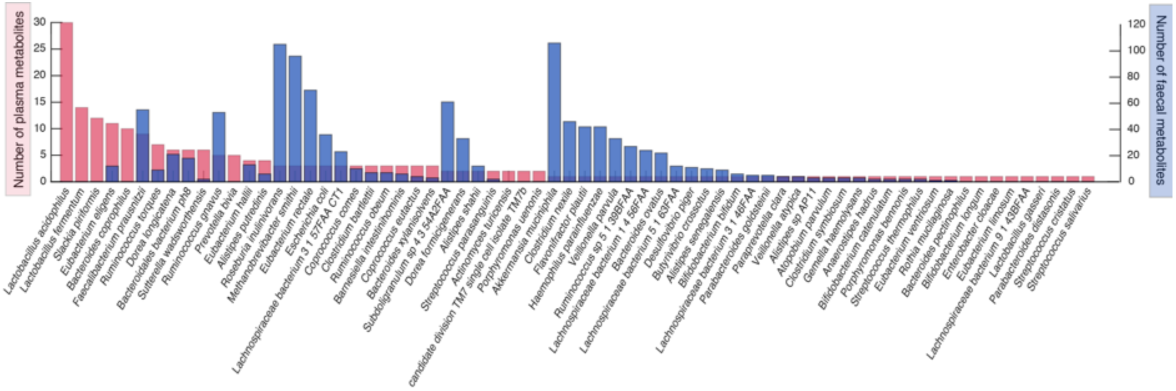


Supplementary Figure S4: Number of blood metabolites significantly associated with microbial species (red bars; left axis). The overlapping blue bars represent the number of faecal metabolites significantly associated with the same species (right axis).

Supplementary Table S1: Population statistics for the individuals in the study dataset.

| | **Value** |
|---|---|
| Age | 64.96 ± 7.78 year |
| Sex (F/M) | 965/39 (96.1/3.9%) |
| MZ/DZ/Singletons | 322/402/280 |
| BMI | 26.17 ± 4.71 kg/m2 |
| Total Fat (%) | 40.00 ± 6.22 |
| Visceral Fat (%) | 38.99 ± 8.98 |

Supplementary Table S2: Number of species and microbial metabolic pathways which are associated to at least one metabolite in faeces/blood. Shared indicates the number of species and pathways, that are associated to at least one metabolite in faeces and in blood.

| | **Faecal-specific** | **Blood-specific** | **Shared** |
|---|---|---|---|
| Microbial species | 112 | 21 | 59 |
| Microbial metabolic pathways | 41 | 14 | 305 |
